# Bonferroni’s correction, not Tukey’s, should be used to control the total number of false positives when making multiple pairwise comparisons in experiments with few replicates

**DOI:** 10.1101/2025.05.01.651705

**Authors:** Adam Zweifach

**Affiliations:** The Department of Molecular and Cell Biology University of Connecticut at Storrs 91 N. Eagleville Rd. Storrs, CT 06269-3125

## Abstract

Statistical tests can be used to help determine whether experimental manipulations produce effects. In tests of means, when more than two groups are compared the total number of Type 1 errors (false positive results) increases unless a correction is used. Tukey’s test is thought to offer good control of the false positive rate and high statistical power when all pairwise comparisons are made. However, the number of replicates in laboratory experiments is often quite low, and small sample sizes can undermine assumptions underlying statistical methods. I used simulations to investigate how well Tukey’s test controls the total number of false positives when there are 3-6 experimental groups and 2-6 experimental replicates, conditions that span the range of typical values, and found that it generates too many. I investigated 11 other approaches to controlling false positives and found that none is as effective as the simple Bonferroni correction or offers much more power. I conclude that researchers should not make all pairwise comparisons using Tukey’s test but instead use Bonferroni’s correction on a limited number of pre-selected comparisons.

## Introduction

Statistical testing of experimental data is a key analysis step at multiple points during the process that leads to discovery of new therapeutic targets, lead compounds, chemical probes, and drugs. Statistical testing of sample means generates p values that indicate the probability of observing data with means at least as different as those obtained if the means of the underlying populations are identical (1–3). Each statistical comparison carries a risk of producing a Type 1 error, or false positive-a p value below the threshold selected for statistical significance-even if there is no true difference. When more than two means are compared, the overall number of false positives increases above the intended significance threshold (called α and by convention usually set at 5%) unless a correction is applied. Several approaches for controlling false positives in the context of multiple comparisons have been developed (4,5). While many can be applied directly to the results of a series of t tests, some advocate performing analysis of variance (ANOVA) first and only proceeding to pairwise comparisons if the ANOVA yields a significant p value as an additional safeguard against false positives (see e.g., (5)), although this approach is not universally accepted because it may unnecessarily decrease statistical power (6). Power, the ability to detect true differences, is discussed further below. However, a key benefit of performing ANOVA first is that models that include treatment together with experimental replicate as a blocking factor (two-factor ANOVA) can be used and comparisons between conditions made using the model’s results for treatment, decreasing the effects of between-replicates variability and creating an extremely effective analysis strategy (7,8).

Approaches for controlling false positives differ in the kind of error rate they seek to manage. Some limit the probability of obtaining one or more false positives among all significant findings-a measure called the false discovery rate (FDR) - which is a relatively lenient standard. Others limit the probability that testing a data set will result in one or more false positives, which is known as the family-wise error rate (FWER) and is a stricter criterion (9). However, controlling the FWER does not necessarily control the total false positive rate (FPR), because while the fraction of data sets generating false positives is controlled, in any given set or “family” of tests more than one false positive may be generated. I suspect that most researchers are most interested in controlling the overall FPR, not the FDR or the FWER. Think of a case in which 20 hits from a screen are being retested. Most researchers would expect that appropriate control of false positives means that averaged over many trials they should expect one hit to demonstrate statistically significant activity if in fact none was active, not that in only one of 20 such trials might several compounds appear active.

The simplest strategy for controlling the FWER is Bonferroni’s correction, which involves adjusting α by dividing the level chosen for significance by the total number of comparisons. For example, conducting 10 comparisons at an α of 0.05 would require using a significance threshold of p < 0.005 per test. However, Bonferroni’s correction reduces statistical power in a manner that increases linearly with the number of comparisons made. Power is higher when differences in means are larger, experimental variability (noise) is smaller, sample size (number of replicates) is larger, or the significance threshold is less stringent (10). To control the FWER while maximizing power, Dunnett’s test is commonly recommended when multiple experimental groups are compared to a single control group. Tukey’s test (Tukey’s HSD) is frequently employed when making all possible pairwise comparisons. Other sophisticated methods, such as “step-up” or “step-down” modifications of Bonferroni’s procedure, have been developed in an attempt to enhance power while still controlling error rates (see (4) for an overview). Much of the work that underlies the modern discovery process involves well-controlled laboratory experiments. Experiments typically involve relatively small sample sizes; performing 3 replicates is common, 2 replicates are not unheard of, and more than 6 replicates are rare. Concerns that small sample sizes can compromise statistical tests have prompted evaluation of the performance of tests for 2 sample means (t tests), showing that these largely work as expected (11,12), but to my knowledge the effects of very small sample sizes when multiple means are compared has not been investigated. I examined how well procedures designed to control the FWER or FDR perform at controlling the overall FPR in experiments with 2-6 replicates and found that Dunnett’s test effectively maintained the FPR in comparisons of multiple treatment groups to a single control, but when multiple pairwise comparisons were made, Tukey’s test often generated a higher-than-desired FPR. Evaluation of alternative procedures revealed that none provided superior control of the FPR or significantly greater power than the Bonferroni method.

## Methods

All simulation were performed using R software (13) running in the R studio IDE. Packages used include multcomp (14) and agricolae (15). To investigate the FPR, two-factor ANOVA was performed on simulated data from 3-6 treatment groups in 2-6 replicates, with the factors being “treatment” and “replicate” (statisticians would call this a “block”, but that term can be confusing to biologists), mimicking the most effective way to analyze more than 2 group means in replicated experimental data (7,8). The mean of each of the groups was set to 100, the standard deviation (SD) of each group was set to 5, and shared variation with an SD of 20 was then added. As mentioned above, although none of the tests I performed require it, I only compared individual group means if the p value of the ANOVA for “treatment” was < 0.05. Although when using one-way ANOVA this is not required, when using two-factor ANOVA it allows effects due to treatment to be isolated, mimicking the most effective analysis strategy. It also ensures that the FWER will be <5% and reduces the overall computational burden by limiting the number of iterations in which pairwise comparisons needed to be calculated. The total number of p values < 0.05 when comparing groups to a single control was summed to generate the overall FPR for Dunnett’s test, and for all pairwise comparisons for other tests. 100,000 simulations were performed to estimate the FPR, except in a preliminary screen conducted to eliminate computationally-intensive methods that failed to control the FPR well that used only 1000 iterations. Estimates of the error with which the FPR is measured after 100,000 iterations ranged from 1% CV for the omnibus ANOVA test to ∼ 3% for the BY procedure which has a very low mean FPR (See Supplemental Table 1). To estimate power, the mean of one group was varied from 2.5 to 50 and the mean of 2-5 others was set to 100. Independent variation was set to 5% and shared variation to 20%. 10,000 simulations were performed, and for each, the number of p values < 0.05 was measured and converted to power by dividing by the total number of p values < 0.05 expected if all real differences were found. An R script is included as supplementary information that will allow any of the calculations in this work to be made.

## Results

### Dunnett’s and Bonferroni’s procedures control the FPR well when sample sizes are small, but Tukey’s test does not

I first examined the FPR following two-factor ANOVA, modeling “treatment” as one factor and “replicate” as the second. If the p value for the ANOVA was < 0.05, I compared all experimental conditions to a single “control” with or without Dunnett’s correction (Figure 1). As mentioned previously, I chose two-factor ANOVA because it helps mitigate the impact of between-run (or between-block) variability on power and is applicable whether samples are independent or matched (7,8). Performing Dunnett’s test only when the ANOVA p value was < 0.05 also ensures that the FWER will not exceed 5%. Simulations were performed for 3-6 experimental groups, each with 2-6 replicates. The total number of p-values < 0.05 was used to estimate the FPR. When only two replicates were simulated, Dunnett’s test resulted in a slightly elevated FPR (∼6.25%) regardless of group number. However, if there were three or more replicates the FPR remained at or below 5%.

**Figure 1:**
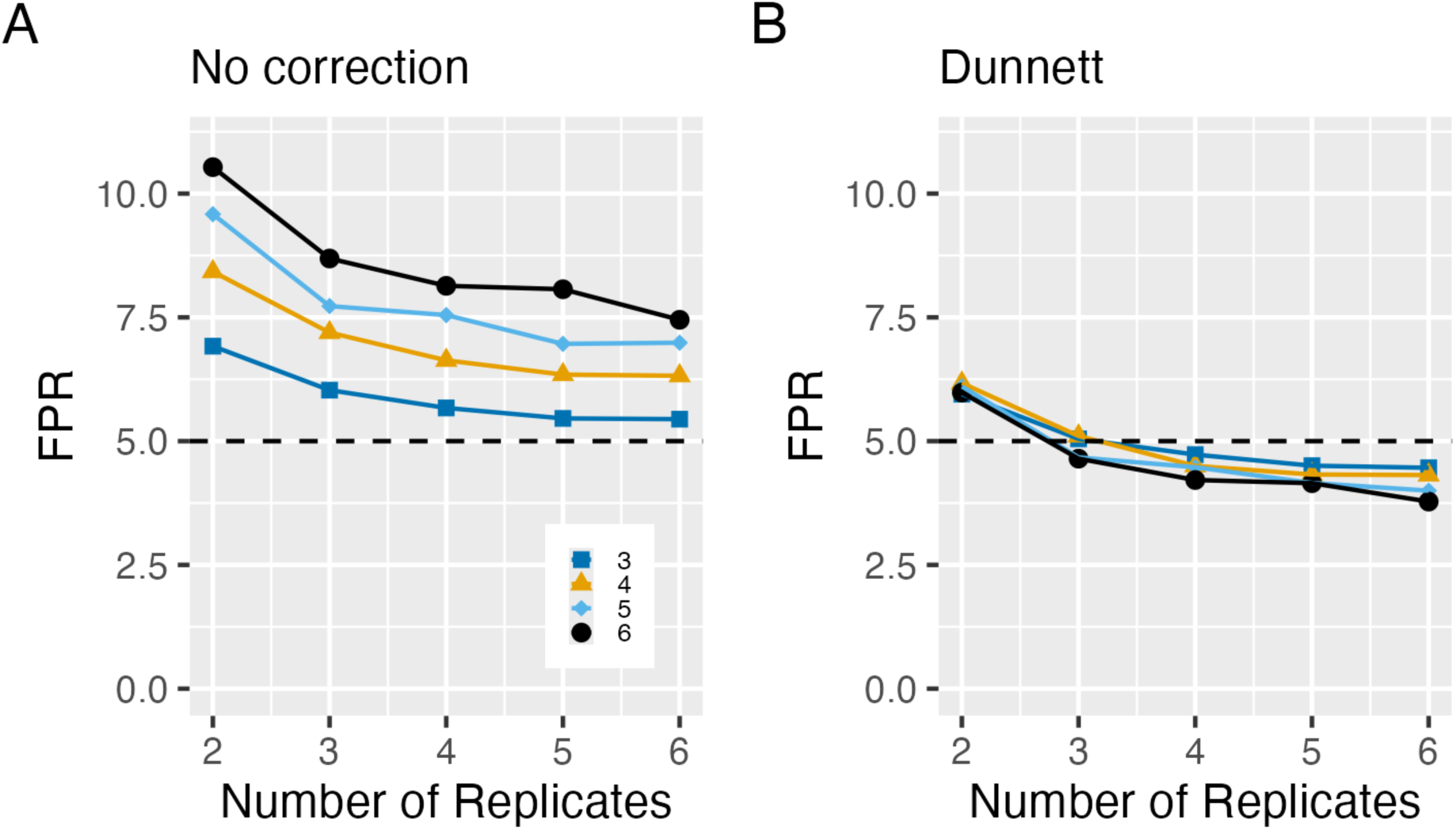
Dunnett’s test controls the FPR when treatment groups are compared to a control. Total rate of false positives (FPR) with no correction (A) or Dunnett’s correction (B). Two-factor ANOVA with one factor being group and the other replicate was performed with the means of 3 (dark blue squares), 4 (mustard triangles), 5 (light blue diamonds) and 6 (black circles) groups all set to 100, independent variation of 5%, and shared variation of 20%, for 2-6 simulated replicates. If the p value of the ANOVA result for “treatment” was < 0.05, comparisons between group 1 and all other groups were made. The total number of p values < 0.05 in 100,000 simulations was used to determine the FPR.

I then conducted a similar analysis making all pairwise comparisons. If the omnibus ANOVA test for treatment was significant (p < 0.05), I summed the number of p-values < 0.05 for pairwise comparisons calculated without correction, with Bonferroni’s correction, or with Tukey’s correction (Figure 2). Bonferroni’s procedure consistently controlled the FPR at 5% across all conditions, but Tukey’s test did not. The highest FPR values for Tukey’s procedure occurred when there were only two replicates and six groups. The FPR decreased as the number of replicates increased or the number of groups decreased. I found that when there are 3 replicates, the FPR reached a maximum at 6-7 treatment groups and then stayed at a constant elevated level as additional groups were added (See Supplemental Figure 1. I confirmed the elevated FPR with Tukey’s test using three different implementations of the procedure in R: glht() from the multcomp package, TukeyHSD() from stats, and pairs() from emmeans (not shown), although I cannot be certain that these represent genuinely independent implementations of the test.

**Figure 2.**
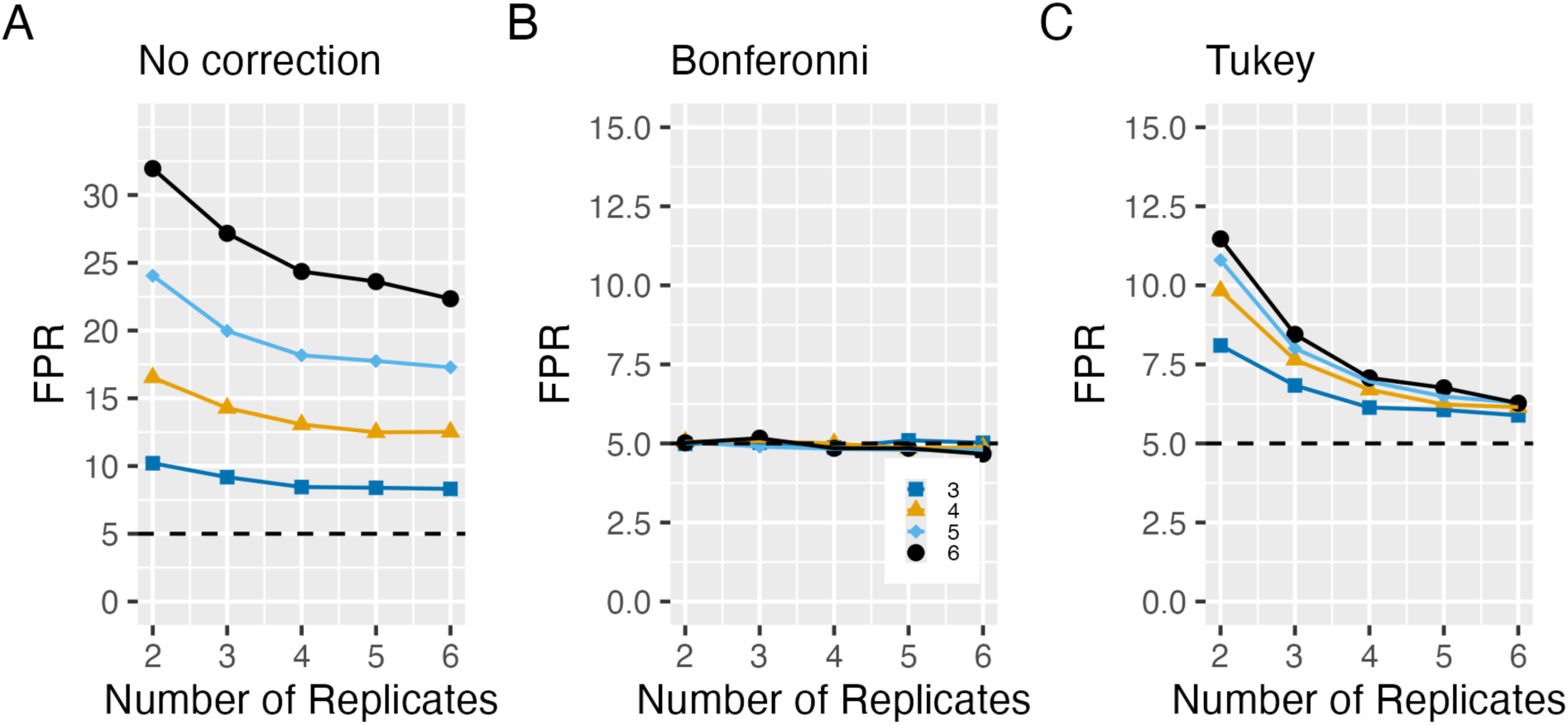
Tukey’s test does not control the FPR well when all pairwise comparisons are made. **FPR** with no correction (A), Bonferroni’s correction (B) or Tukey’s correction (C) when all possible pairwise connections are made with the means of 3 (dark blue squares), 4 (mustard triangles), 5 (light blue diamonds) and 6 (black circles) groups all set to 100, independent variation of 5%, and shared variation of 20%, for 2-6 simulated replicates. The total number of p values < 0.05 in 100,000 simulations was used to determine the FPR.

### Testing the efficacy of other strategies that control the FPR

I next tested 11 other procedures for controlling either the FPR, all of which are available in R and could be applied after two-factor ANOVA. These included the Benjamini-Hochberg (BH), Benjamini-Yekutieli (BY), Holm, Hochberg, Shaffer, Sidak, and Westfall procedures from package multcomp, and the Duncan, REGW, Scheffé, and SNK procedures from package Agricolae (see references (16,17)). Methods supported by the package multcomp make use of the full ANOVA object, while those from package agricolae use a different strategy involving an estimate of the variance associated with the factor “treatment” and thus may not be fully comparable to the multcomp methods. Initial comparisons were based on 1,000 simulations to limit computation time. Of the methods examined, the BY, Hochberg, Holm, Hommel, Scheffé, and Shaffer procedures reduced the FPR more effectively than Tukey’s test. I then ran 100,000 simulations for the most promising methods (Figure 3). Among these, BY consistently kept the FPR well below 5%, Bonferroni maintained it at 5%, and Holm kept it at or below 6%.

**Figure 3.**
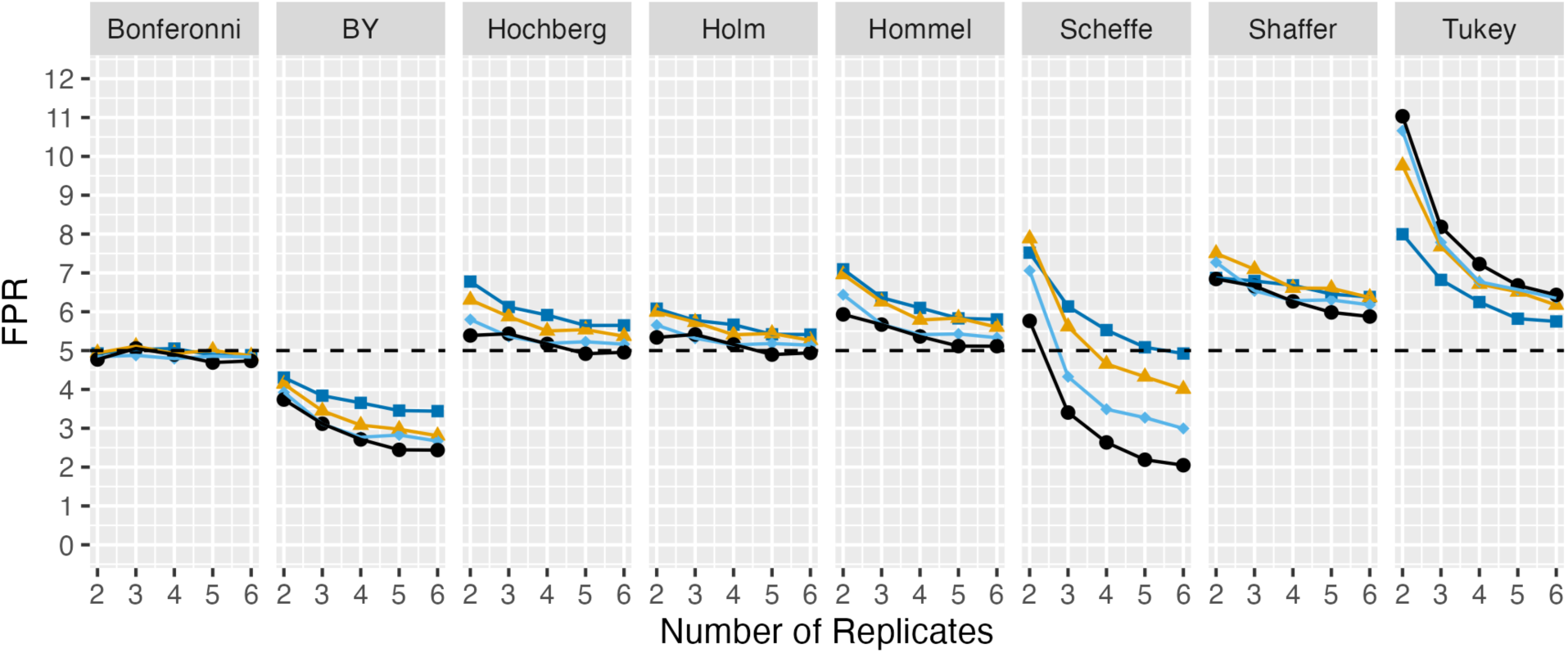
FPR with a selection of other control strategies when all pairwise comparisons are made. 11 approaches to controlling the FPR were tested in an initial round using 1000 simulations (Supplemental Figure 2). 7 were selected for comparison to Bonferroni’s and Tukey’s approaches using 100,000 simulations. Colors and symbols are as in Figure 1 and 2.

Finally, I compared the statistical power of the BY, Holm, and Bonferroni procedures to Tukey’s test. I simulated experiments in which the mean of one group differed from those of 2-5 other groups using 3 replicates per group. Power was calculated as the number of significant p-values < 0.05 divided by (n_groups_ − 1), the maximum number of true positives expected if all real differences were detected. Consistent with their respective impacts on the FPR, Tukey’s test provided the highest power, and BY the lowest. The Holm procedure had higher power than Bonferroni, but the advantage decreased as the number of groups increased.

## Discussion

Using Tukey’s test, the FPR following two-factor ANOVA can exceed 8% when three replicates-perhaps the most common value reported in laboratory experiments-are performed, even if the FWER is maintained at 5% by performing pairwise testing only if the p value for the ANOVA factor treatment is < 0.05. I suspect that most researchers who use Tukey’s test wish to control the overall FPR, not the FWER. A p-value is intended to convey the probability of observing data differing as much as those collected assuming the null hypothesis, that means are the same, is true. If a p-value < 0.05 does not mean that there is a 5% chance of such data under the null, but rather some other chance, its already limited interpretive value is further diminished.

I simulated two-factor ANOVA because it is the best choice for analyzing replicated experiments. If there is variation between replicates, two-factor ANOVA minimizes its effects and results in high power when post-hoc comparisons are made using the results for the “treatment” factor. If there is no or minimal variation between replicates, two-way ANOVA gives results that differ only slightly from one-way ANOVA. However, I did confirm that the effects of the various procedures on the FPR that I observed using two-factor ANOVA held true following one-way ANOVA (Supplemental Figure 3). I also had ChatGPT generate a completely independent simulation of both the FWER and FPR when one way ANOVA is followed by Bonferroni’s procedure or Tukey’s test for 6 experimental conditions and 2-10 experimental replicates (Supplemental Figure 4). The results confirm that Tukey’s test maintains the FWER but that the FPR is elevated.

Among the methods tested for controlling false positives when making pairwise comparisons, none performed as well at controlling the FPR as the Bonferroni correction or offered a substantial gain in power. This makes Bonferroni the best option. However, it is worth noting that using Bonferroni’s correction does reduce power compared to using Tukey’s test (Figure 4). While power can be increased by adding replicates, this requires additional time and resources. Fortunately, power can also be enhanced simply by reducing the total number of comparisons made. Of course, this requires pre-specifying a subset of comparisons of interest before the experiment begins, an approach that has the advantage of preventing “HARKing” (Hypothesizing After the Results are Known) and data dredging (18,19). Importantly, focusing on preselected contrasts does not mean sacrificing a valuable statistical tool, as making all pairwise comparisons is rarely a good idea. Pairwise comparisons that are expected to yield non-significant results are useless and should not be made, because when a test yields a p-value above the chosen significance threshold, the correct interpretation is simply that there is no evidence that the population means differ, not that they are the same. Moreover, two major classes of experiments that are frequently analyzed using all-pairs comparisons followed by Tukey’s test are better handled in other ways. The first class includes factorial experiments where the effect of one variable (e.g., presence vs. absence of drugs, or genetic manipulations) is assessed in the context of a second variable that also has two or more levels. These studies are best analyzed using a two-factor ANOVA with an interaction term. This directly tests whether the effect of one factor depends on the level of the other, a question that pairwise comparisons cannot answer. The second class includes dose-response or time-course experiments, which may also be conducted across different conditions such as genotypes or drug types. Rather than performing all-pairs comparisons at each time point or dose, more appropriate approaches include linear regression (20), which can be performed on raw or transformed data, mixed models, analysis of theory-derived parameters (e.g., IC₅₀, K_d_, V_max_, time constants, etc.), or statistical tests performed on summary measures like area under the curve (AUC). These approaches incorporate the structure of time and/or dose into the analysis, making use of important information that Tukey’s test ignores and gaining power as a result. In fact, in these contexts, using all-pairs comparisons generally results in scientifically meaningless conclusions, because whether an observed difference between specific time points, conditions, or doses is statistically significant depends as much on sample size and variability as on the size of the effect. Note that methods like t tests and ANOVA typically assume that sample data are normally distributed. This may not always be the case, particularly for potency data, which often follow a log-normal distribution (see e.g.(21)). It is important researchers have some sense of how data are distributed to ensure that appropriate methods are being used.

**Figure 4.**
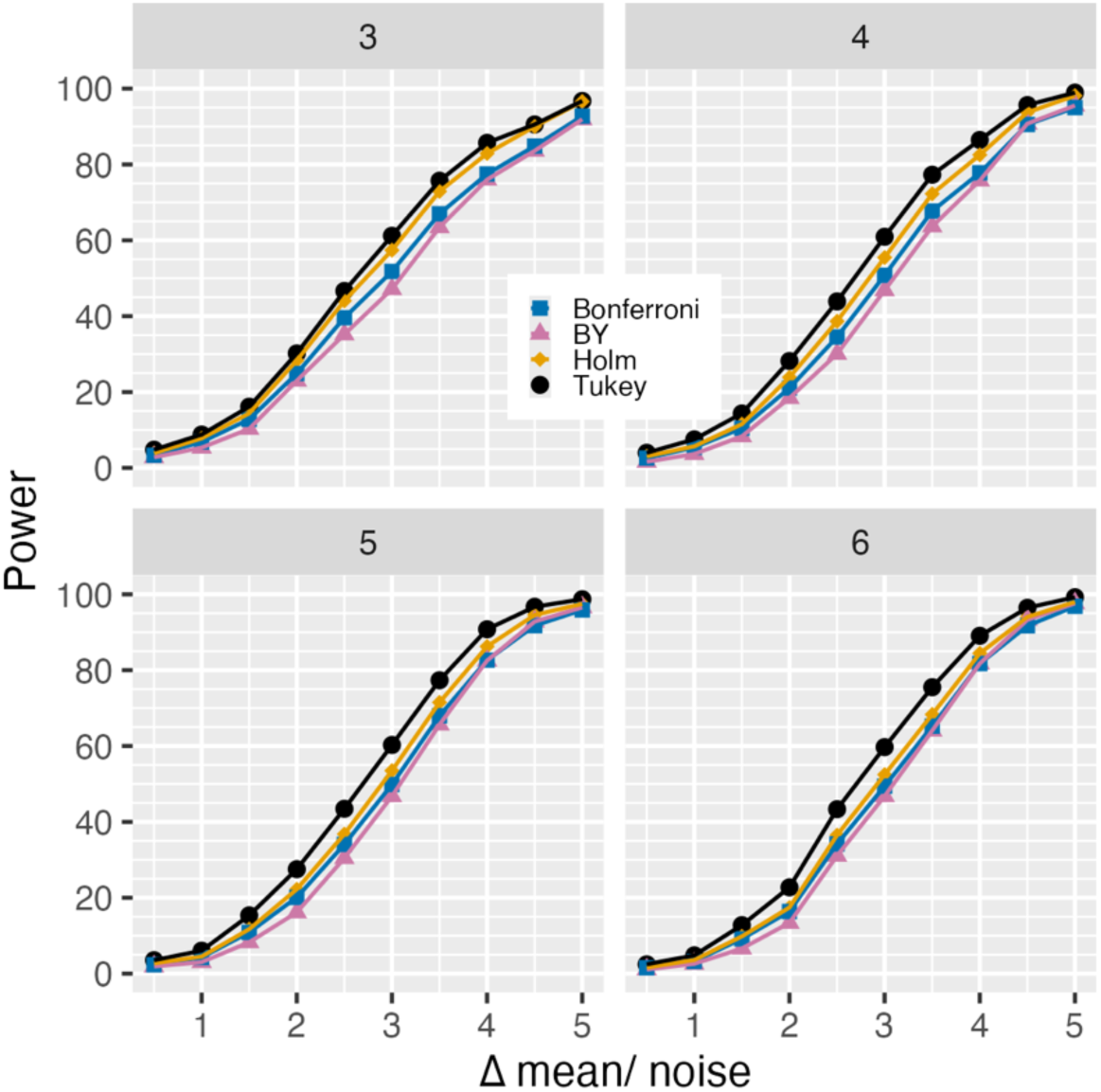
Power of Bonferroni, BY, Holm and Tukey procedures when making pairwise comparisons. Plots of power of the Bonferroni (blue squares), BY (mauve triangles), Holm (mustard diamonds) and Tukey procedures (black circles) following two-factor ANOVA. Power was calculated by dividing the total number of p values < 0.05 in each simulation by the total number of possible differences, n-1, where n is the number of groups. The x axis of each plot is the difference in mean of the first group divided by independent variation, a convenient way of normalizing effect sizes. 10,000 simulations were performed with the mean of one group set to values ranging from 2.5 to 50 and the mean of 2-5 others set to 100, with independent variation of 5% and shared variation of 20%. The total number of groups is indicated in each panel’s banner.

When experiments of the two kinds discussed above are analyzed using methods more appropriate to their design, few scenarios remain in which all-pairs comparison is the best option. Of relevance to the readership of *SLAS Discovery*, the most appropriate context for all-pairs comparisons may be exploratory early-stage research. There, though, the goal is hypothesis generation rather than formal testing. A strong argument can be made that no correction for multiple comparisons should be applied when results are intended to guide future experiments, as those will reveal which effects are genuine, and correction for multiple comparisons might decrease power to find interesting candidates to pursue.

## Supplemental information

**Supplementary Figure 1:**
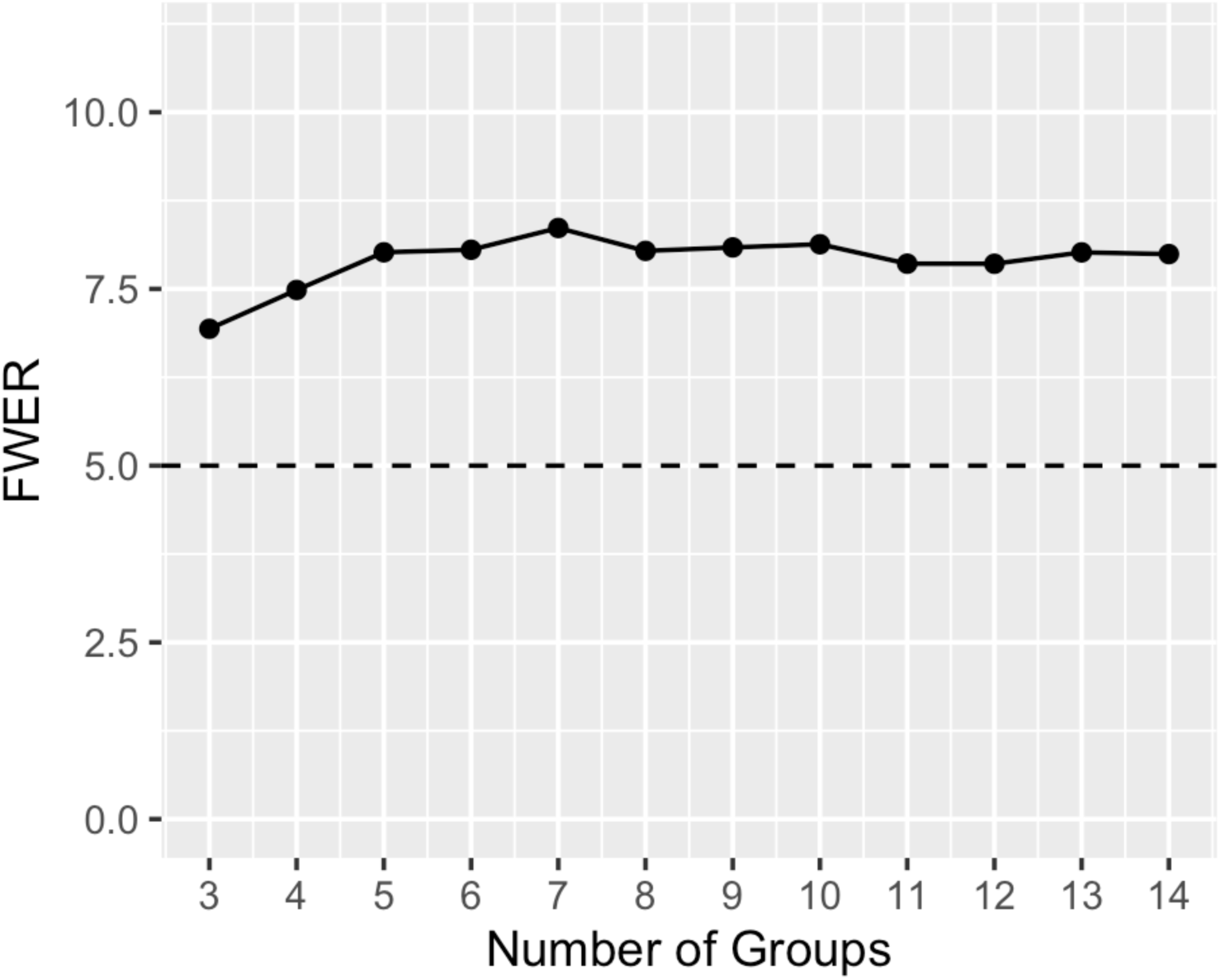
FPR of Tukey’s test with 3 replicates and from 3-15 sample groups.

**Supplementary Figure 2:**
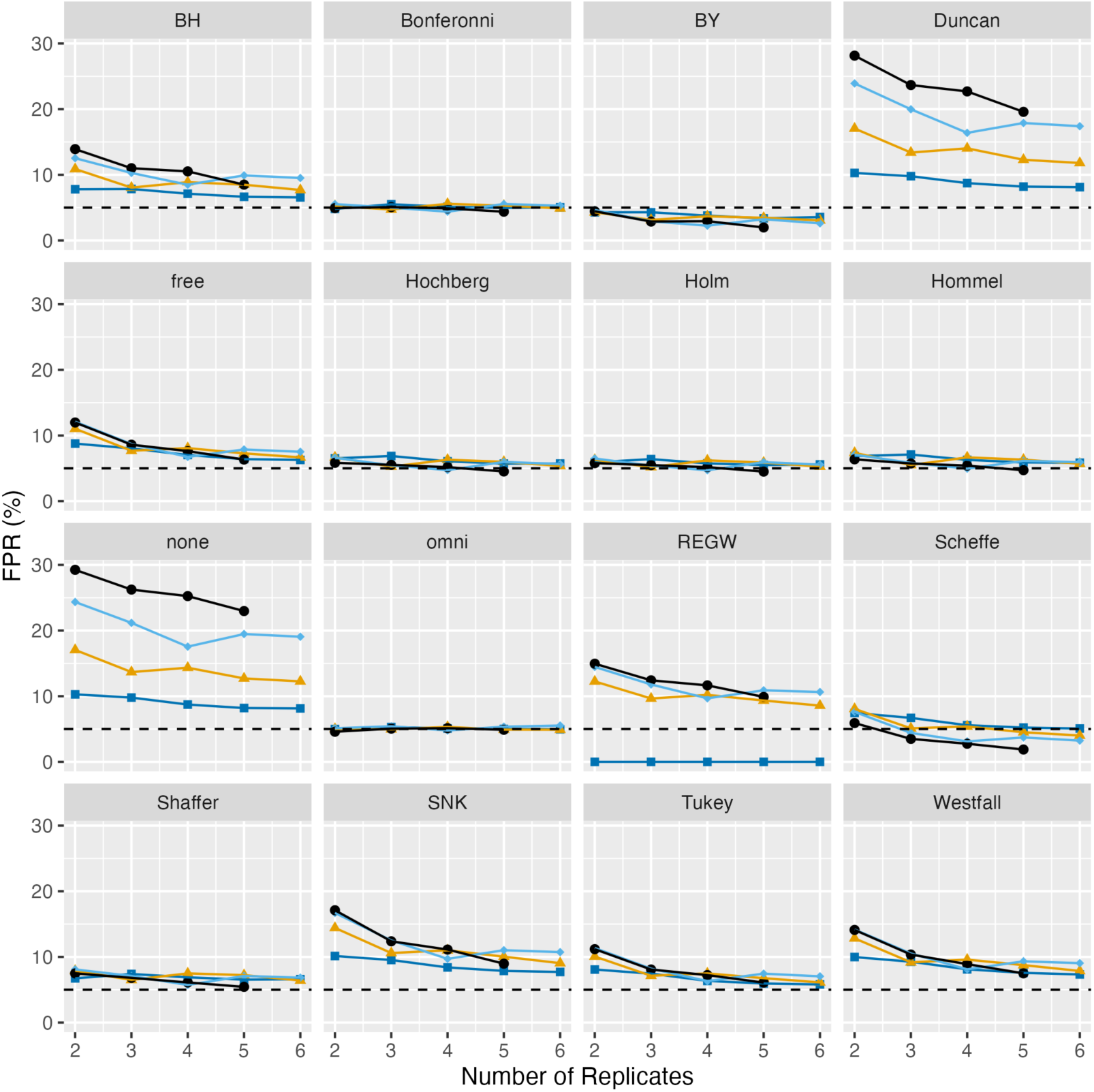
Screening other strategies for controlling the FPR with 10000 iterations prior to select a subset for more detailed analysis.

**Supplementary Figure 3:**
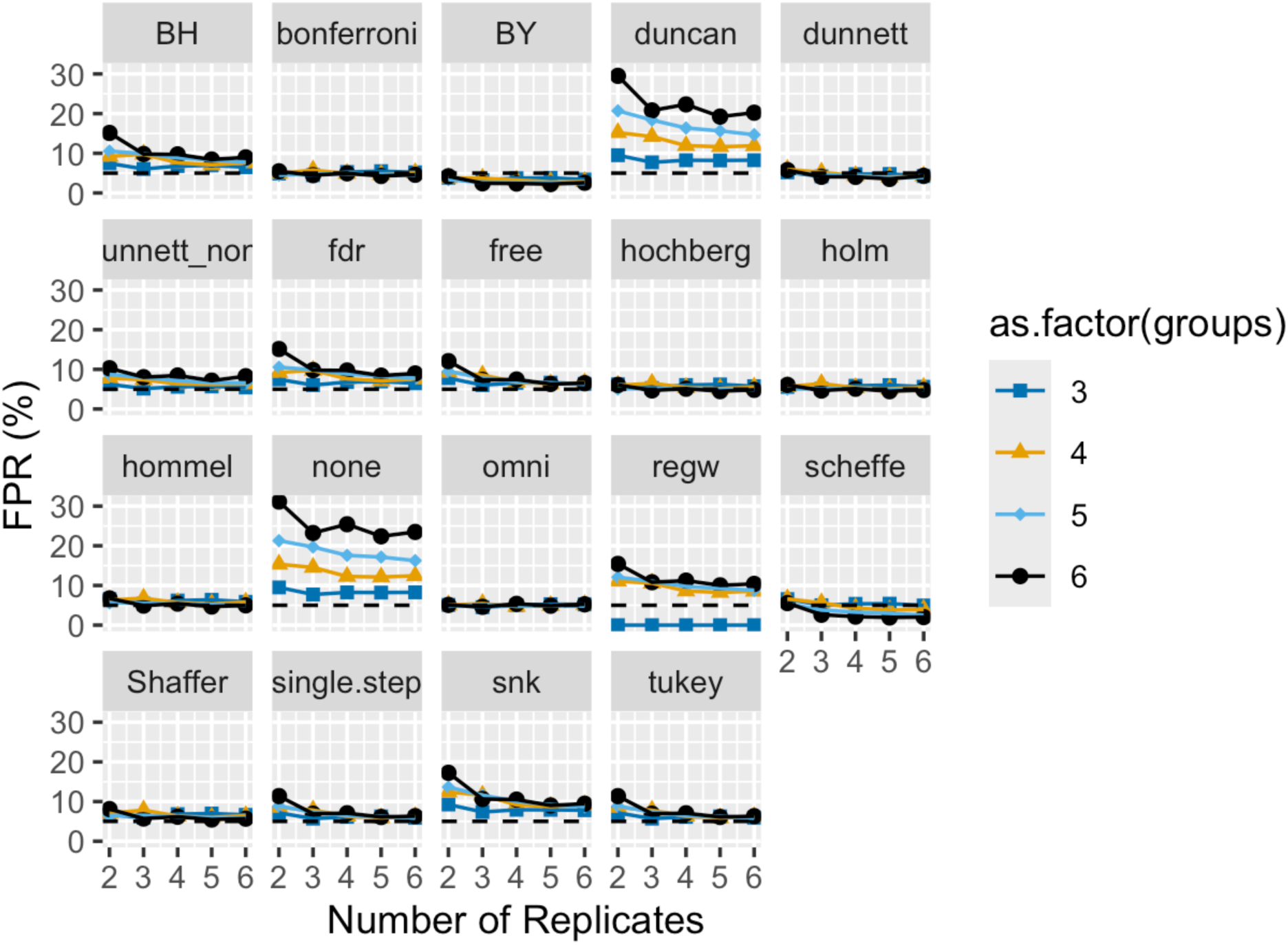
FPR following one-way ANOVA assessed with 10,000 iterations.

**Supplementary Figure 4:**
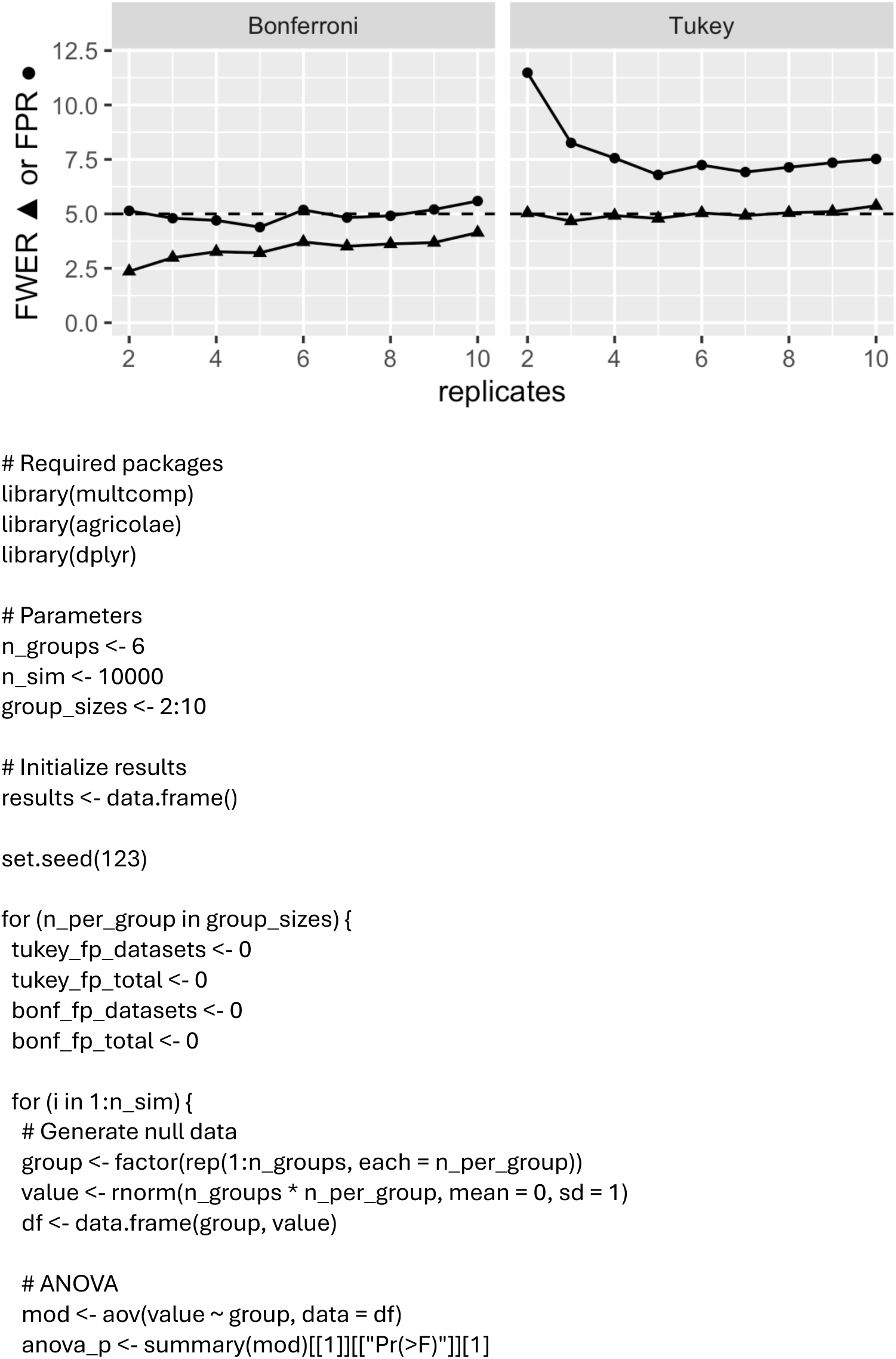
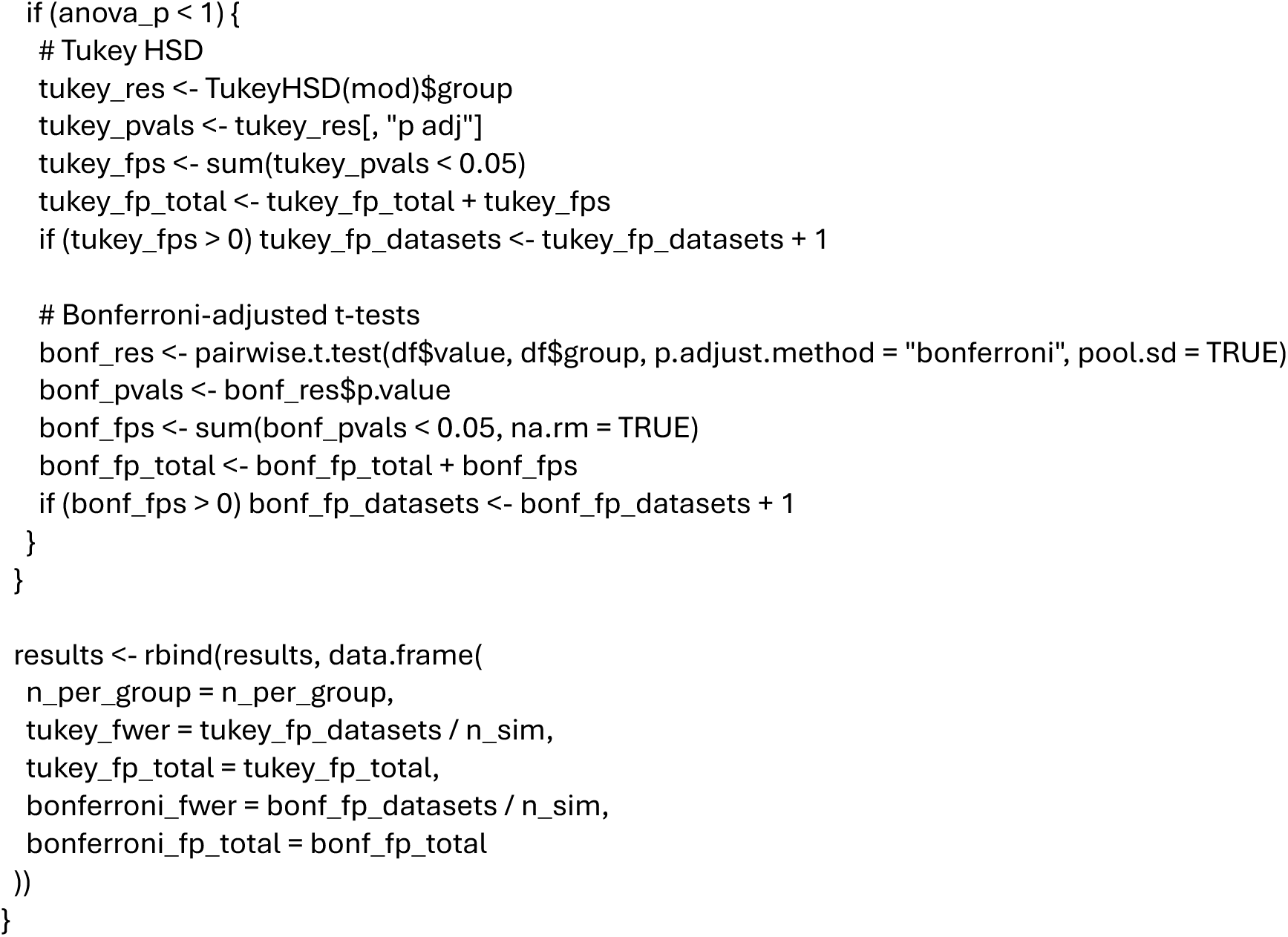
Independent simulation of the FWER and FPER following one-way ANOVA. ChatGPT was prompted to generate a simulation with no user input. 6 experimental groups were simulated in 10,000 iterations. Code is included below.

**Supplemental Table:**
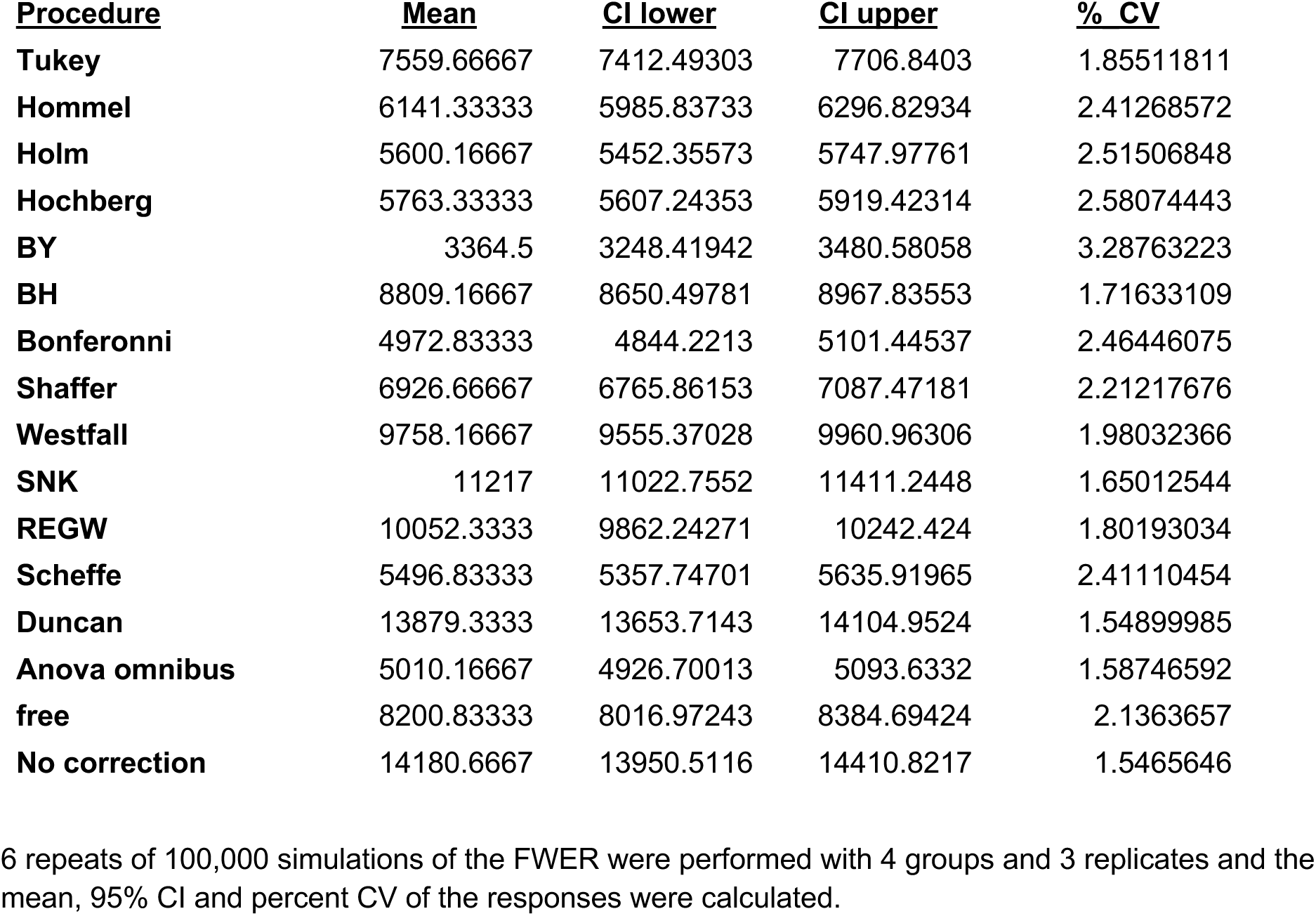
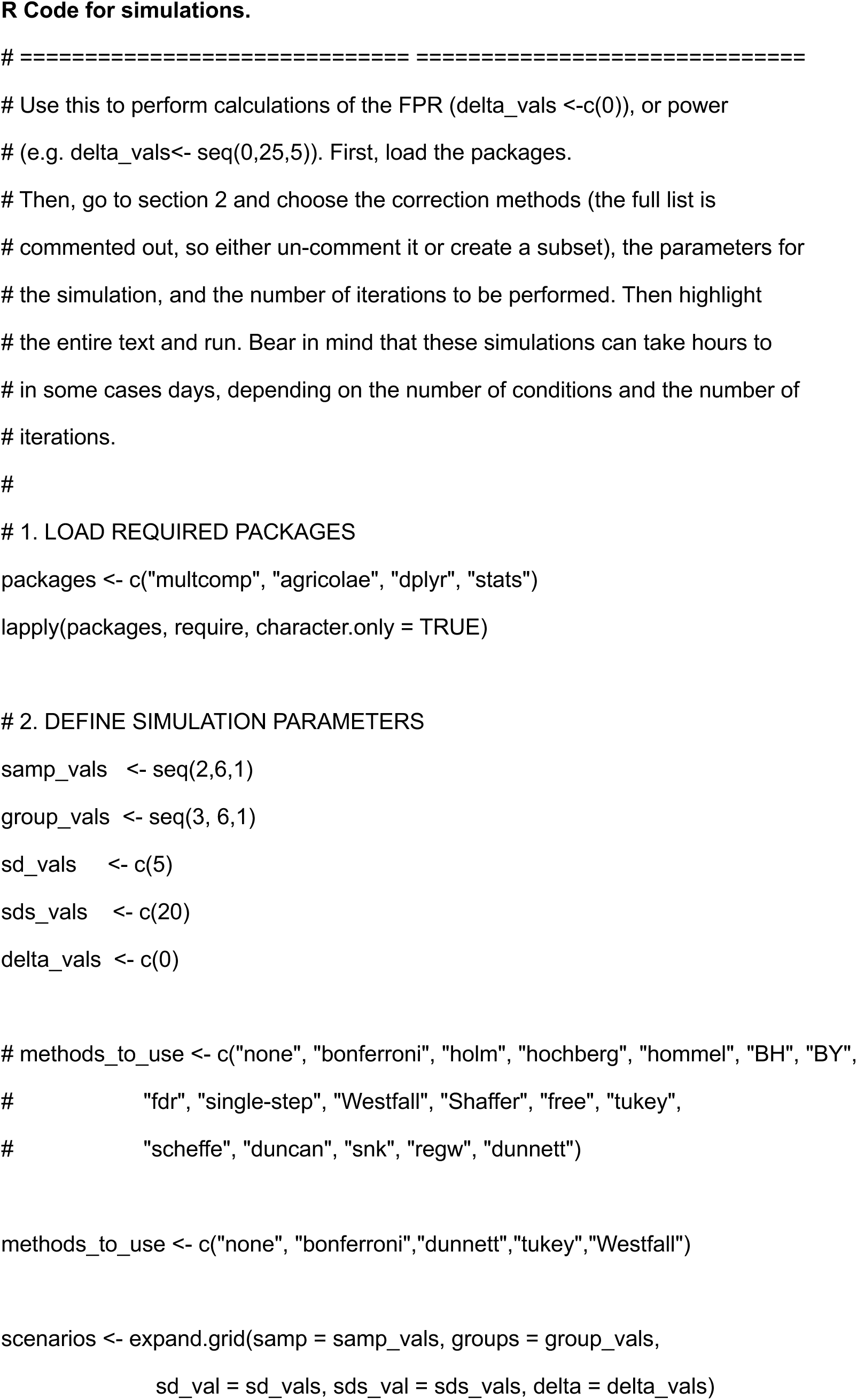
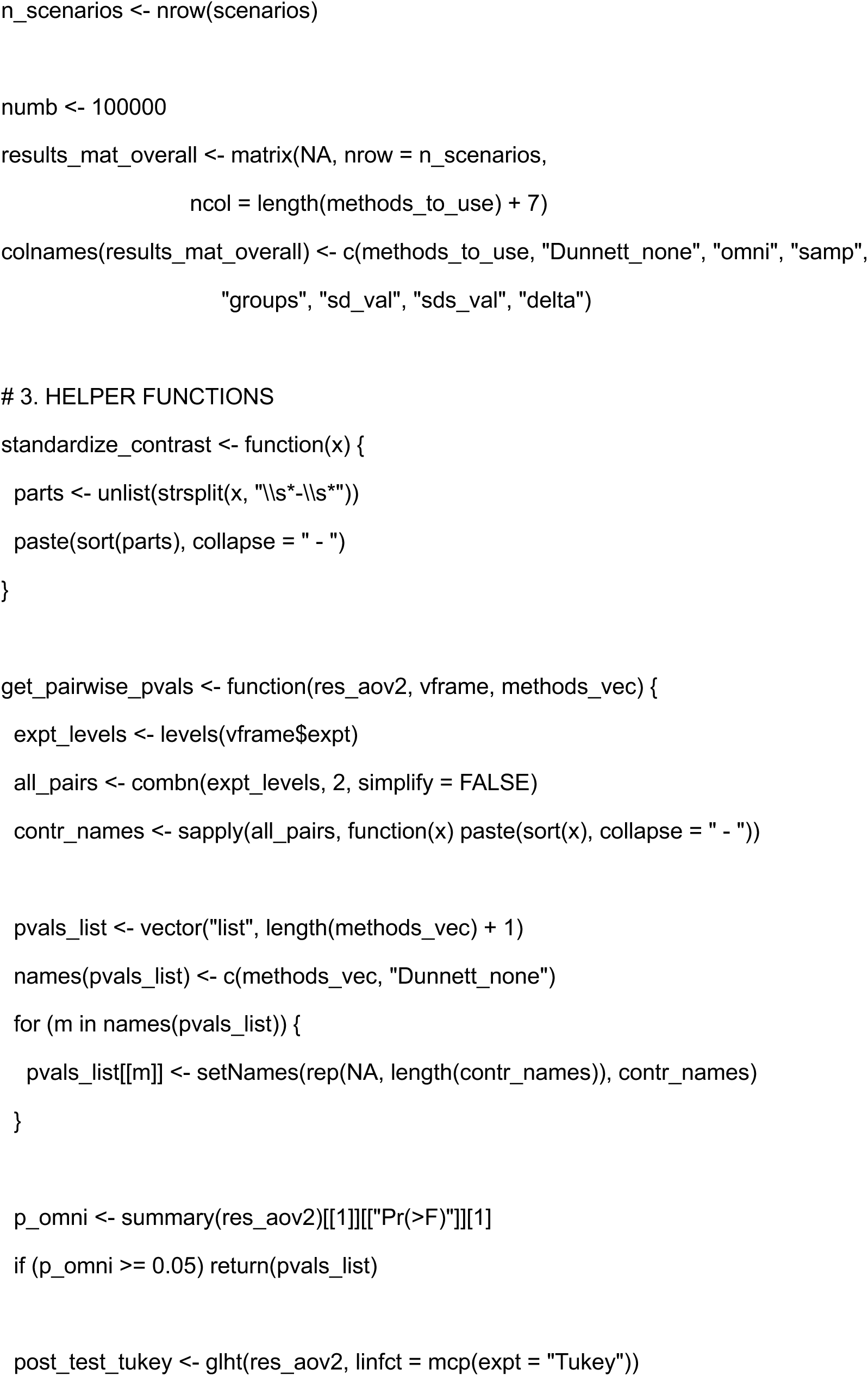
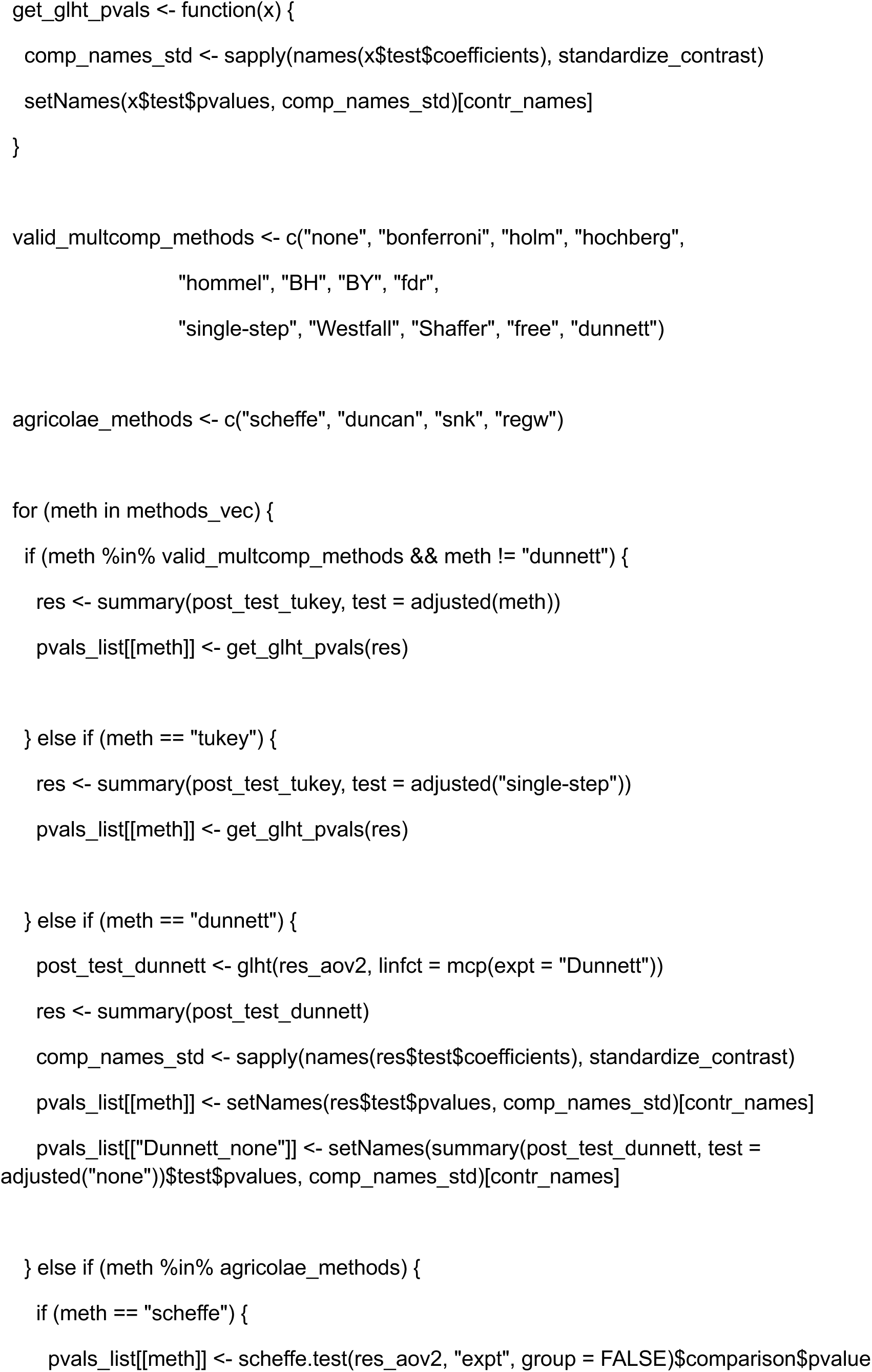
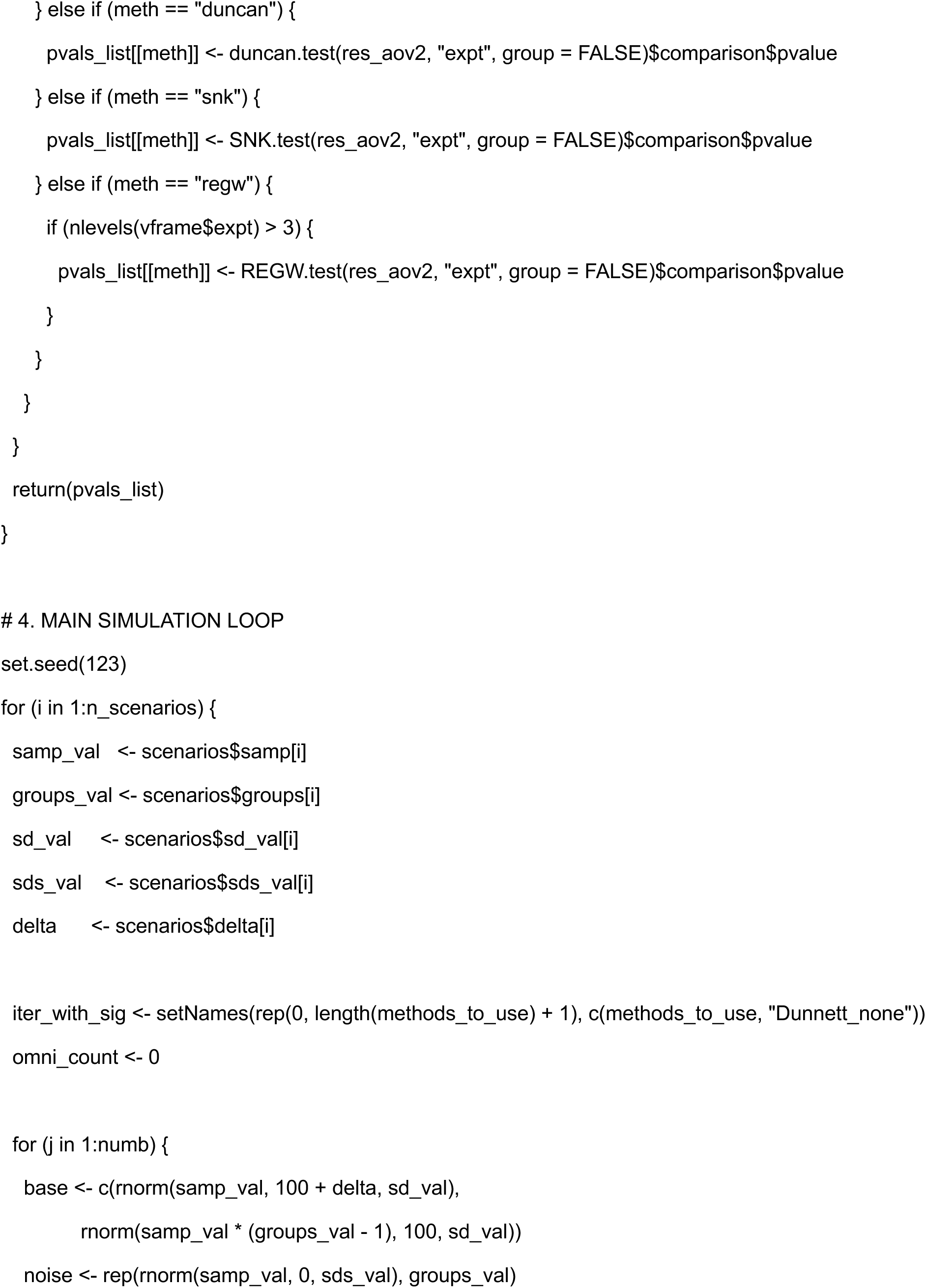
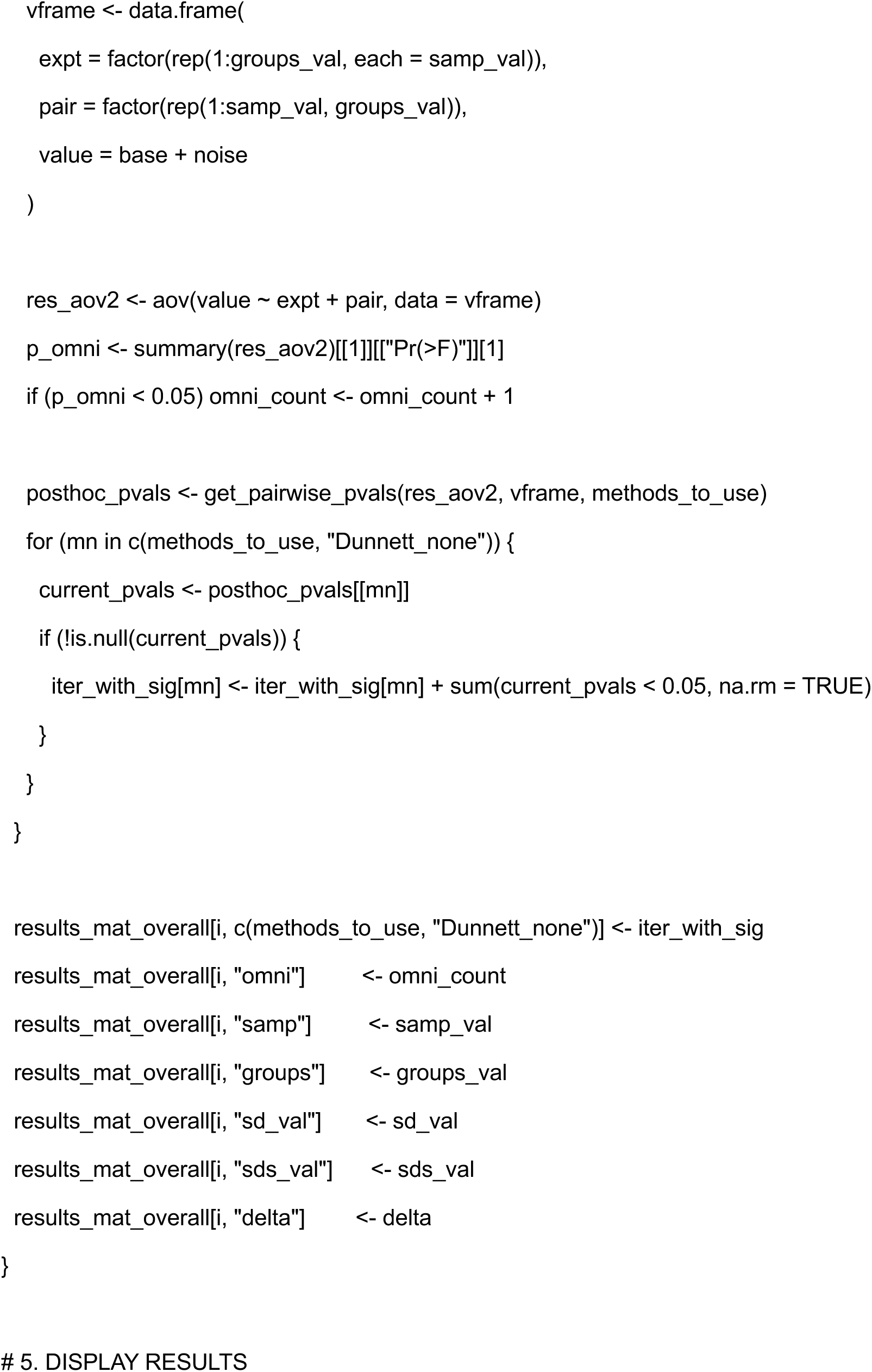
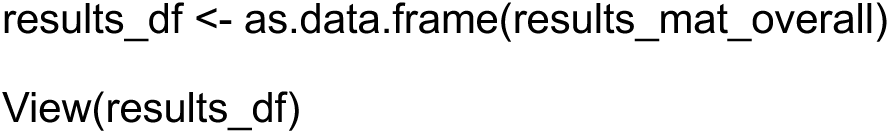
Errors associated with estimates of the FPR.

## References

1. Davis RB, Mukamal KJ. Hypothesis Testing. Circulation. 2006 Sep 5;114(10):1078– 82.

2. Emmert-Streib F, Dehmer M. Understanding Statistical Hypothesis Testing: The Logic of Statistical Inference. Machine Learning and Knowledge Extraction. 2019 Sep;1(3):945–61.

3. Szucs D, Ioannidis JPA. When Null Hypothesis Significance Testing Is Unsuitable for Research: A Reassessment. Frontiers in Human Neuroscience [Internet]. 2017 Aug 3 [cited 2019 Apr 10];11. Available from: https://www.ncbi.nlm.nih.gov/pmc/articles/PMC5540883/

4. Midway S, Robertson M, Flinn S, Kaller M. Comparing multiple comparisons: practical guidance for choosing the best multiple comparisons test. PeerJ. 2020;8:e10387.

5. Lee S, Lee DK. What is the proper way to apply the multiple comparison test? Korean J Anesthesiol. 2018 Oct;71(5):353–60.

6. Hothorn LA. The two-step approach—a significant ANOVA F-test before Dunnett’s comparisons against a control—is not recommended. Communications in Statistics - Theory and Methods. 2016 Jun 2;45(11):3332–43.

7. Lew M. Good statistical practice in pharmacology. Problem 2. Br J Pharmacol. 2007 Oct;152(3):299–303.

8. Zweifach A. Samples in many cell-based experiments are matched/paired but taking this into account does not always increase power of statistical tests for differences in means. MBoC. 2024 Jan;35(1):br1.

9. Finner H, Roters M. On the False Discovery Rate and Expected Type I Errors. Biometrical Journal. 2001;43(8):985–1005.

10. Krzywinski M, Altman N. Power and sample size. Nature Methods. 2013 Dec 1;10(12):1139–40.

11. de Winter JCF. Using the Student’s t-test with extremely small sample sizes. Practical Assessment, Research, and Evaluation [Internet]. 2019 [cited 2021 Nov 9];18. Available from: https://scholarworks.umass.edu/pare/vol18/iss1/10/

12. Poncet A, Courvoisier DS, Combescure C, Perneger TV. Normality and Sample Size Do Not Matter for the Selection of an Appropriate Statistical Test for Two-Group Comparisons. Methodology. 2016 Apr;12(2):61–71.

13. R Core Team. R: A Language and Environment for Statistical Computing [Internet]. Vienna, Austria: R Foundation for Statistical Computing; 2020. Available from: https://www.R-project.org/

14. Hothorn T, Bretz F, Westfall P. Simultaneous Inference in General Parametric Models. Biometrical Journal. 2008;50(3):346–63.

15. Mendiburu F de. agricolae: Statistical Procedures for Agricultural Research [Internet]. 2023. Available from: https://CRAN.R-project.org/package=agricolae

16. Kim HY. Statistical notes for clinical researchers: post-hoc multiple comparisons. Restor Dent Endod. 2015 May;40(2):172–6.

17. Benjamini Y, Yekutieli D. The control of the false discovery rate in multiple testing under dependency. The Annals of Statistics. 2001 Aug;29(4):1165–88.

18. Stefan AM, Schönbrodt FD. Big little lies: a compendium and simulation of p-hacking strategies. Royal Society Open Science. 2023 Feb 8;10(2):220346.

19. Head ML, Holman L, Lanfear R, Kahn AT, Jennions MD. The Extent and Consequences of P-Hacking in Science. PLOS Biology. 2015 Mar 13;13(3):e1002106.

20. Lazic SE. Why we should use simpler models if the data allow this: relevance for ANOVA designs in experimental biology. BMC Physiology. 2008;8:16.

21. Elassaiss-Schaap J, Duisters K. Variability in the Log Domain and Limitations to Its Approximation by the Normal Distribution. CPT Pharmacometrics Syst Pharmacol. 2020 May;9(5):245–57.

